# THE RHODOEXPLORER PLATFORM FOR RED ALGAL GENOMICS AND WHOLE GENOME ASSEMBLIES FOR SEVERAL GRACILARIA SPECIES

**DOI:** 10.1101/2023.03.20.533491

**Authors:** Agnieszka P. Lipinska, Stacy A. Krueger-Hadfield, Olivier Godfroy, Simon Dittami, Lígia Ayres-Ostrock, Guido Bonthond, Loraine Brillet-Guéguen, Susana Coelho, Erwan Corre, Guillaume Cossard, Christophe Destombe, Paul Epperlein, Sylvain Faugeron, Elizabeth Ficko-Blean, Jessica Beltrán, Emma Lavaut, Arthur Le Bars, Fabiana Marchi, Stéphane Mauger, Gurvan Michel, Philippe Potin, Delphine Scornet, Erik E. Sotka, Florian Weinberger, Mariana Cabral de Oliveira, Marie-Laure Guillemin, Estela M. Plastino, Myriam Valero

**Affiliations:** Department of Algal Development and Evolution, Max Planck Institute for Biology Tubingen, Tubingen, Germany; Department of Biology, University of Alabama at Birmingham, 1300 University Blvd, Birmingham, AL, 35294; Sorbonne Université, CNRS, UMR 8227, Laboratory of Integrative Biology of Marine Models, Station Biologique de Roscoff, Roscoff, France; Departamento de Botânica, Instituto de Biociências, Universidade de São Paulo, Rua do Matão 277, Cidade Universitária 05508-090, São Paulo, SP, Brasil; Hortimare - Breeding & Propagating Seaweed. Altonstraat 25A 1704 CC Heerhugowaard. The Netherlands; Institute for Chemistry and Biology of the Marine Environment (ICBM), Carl von Ossietzky University Oldenburg, Schleusenstrasse 1, 26382, Wilhelmshaven, Germany; CNRS, Sorbonne Université, FR2424, ABiMS-IFB, Station Biologique, 29680, Roscoff, France; CNRS, Sorbonne Université, Pontificia Universidad Católica de Chile, Universidad Austral de Chile, IRL 3614, Evolutionary Biology and Ecology of Algae, Station Biologique de Roscoff, CS 90074, F-29688 Roscoff, France; Núcleo Milenio MASH, Facultad de Ciencias Biológicas, Pontificia Universidad Católica de Chile, Santiago, Chile; CNRS, Institut Français de Bioinformatique, IFB-core, UMS 3601, Évry, France; Department of Biology, College of Charleston, Charleston SC 29412; GEOMAR Helmholtz-Zentrum für Ozeanforschung, Marine Ecology Division, Düsternbrooker Weg 20, 24105 Kiel, Germany; Núcleo Milenio MASH, Facultad de Ciencias, Instituto de Ciencias Ambientales y Evolutivas, Universidad Austral de Chile, Casilla 567, Valdivia, Chile; Centro FONDAP de Investigación de Ecosistemas Marinos de Altas Latitudes (IDEAL), Valdivia, Chile

**Author notes:** Authors for correspondence: Agnieszka P. Lipinska, Department of Algal Development and Evolution, Max Planck Institute for Developmental Biology, Tuebingen, Germany; Stacy A. Krueger-Hadfield, Department of Biology, University of Alabama at Birmingham, 1300 University Blvd, Birmingham, AL, 35294. Shared first authors (Agnieszka P. Lipinska, Stacy A. Krueger-Hadfield).

**Keywords:** evolution, ecology, omics, ploidy, Rhodophyta

## Abstract

Macroalgal (seaweed) genomic resources are generally lacking as compared to other eukaryotic taxa, and this is particularly true in the red algae (Rhodophyta). Understanding red algal genomes is critical to understanding eukaryotic evolution given that red algal genes are spread across eukaryotic lineages from secondary endosymbiosis and red algae diverged early in the Archaeplastids. The Gracilariales are highly diverse and widely distributed order whose species can serve as ecosystem engineers in intertidal habitats, including several notorious introduced species. The genus *Gracilaria* is cultivated worldwide, in part for its production of agar and other bioactive compounds with downstream pharmaceutical and industrial applications. This genus is also emerging as a model for algal evolutionary ecology. Here, we report new whole genome assemblies for two species (*G. chilensis* and *G. gracilis*), a draft genome assembly of *G. caudata*, and genome annotation of the previously published *G. vermiculophylla* genome. To facilitate accessibility and comparative analysis, we integrated these data in a newly created web-based portal dedicated to red algal genomics (https://rhodoexplorer.sb-roscoff.fr). These genomes will provide a resource for understanding algal biology and, more broadly, eukaryotic evolution.

## SIGNIFICANCE STATEMENT

The Gracilariales are an ecologically and economically important red algal order found throughout the coastal regions of the world. Understanding the biology, ecology, and evolution of species in this order, and that of red algae more broadly, has been hampered by the limited phylogenetic coverage of genomic resources. Here, we present whole genome assemblies and gene annotations for four *Gracilaria* species that will serve as a key resource for algal research on evolution, ecology, biotechnology and aquaculture.

## INTRODUCTION

Red algae (Rhodophyta) represent a lineage of photosynthetic eukaryotes in the Archaeplastids that diverged from green algae around 1700 MYA (Yang et al. 2016). Within the Rhodophyta, the Cyanidiophyceae were the earliest to diverge approximately 1200 MYA, while the Florideophyceae diverged more recently (i.e., 412 MYA; Yang et al. 2016) and constitute the most speciose group (Graham et al. 2016). In this context, the genomic resources currently available (Table S1) represent only a fraction of the evolutionary diversity of red algae, limiting our capacity to reconstruct the evolutionary history of the unique features of this group.

The Florideophyceae have a life cycle in which haploid male and female gametophytes alternate with a diploid tetrasporophyte (Figure S1). After fertilization, the zygote develops on the female into a cystocarp, in which the zygote is mitotically copied. Male gametes (spermatia) and spores are non-motile, and the female gamete (carpogonium) is retained on the female thallus. The cystocarp was thought to have evolved in response to low fertilization success (Searles 1980), but recent work has shown that many males fertilize a single female (Engel et al. 1999, Krueger-Hadfield et al. 2015) and that animal-mediated fertilization can increase reproductive success (Lavaut et al. 2022). Many species have ‘isomorphic’ gametophytes and tetrasporophytes, which are hard to discern without the aid of molecular tools (e.g., sex-linked markers, Martinez et al. 1999; Guillemin et al. 2012; or microsatellites, Krueger-Hadfield et al. 2016).

Here, we focus on four *Gracilaria*^1^ species spanning roughly 170 million years of evolution (Lyra et al. 2021). These species were chosen based on their evolutionary, ecological, and/or economic importance. Species in the genus *Gracilaria* produce agars in their cell wall (Popper et al. 2011), they can be propagated vegetatively, and serve as ecosystem engineers in intertidal zone (Kain and Destombe 1995). The four taxa chosen can be divided into three clades based on their molecular divergence: (i) *G. chilensis* and *G. vermiculophylla*, (ii) *G. caudata*, and (iii) *G. gracilis* (Lyra et al. 2021). *Gracilaria gracilis* and *G. caudata* are evolutionarily more distinct than the phylogenetic group that contains *G. chilensis* and *G. vermiculophylla*. *Gracilaria chilensis* C.J. Bird et al. is an important crop along the Chilean coastline, where it has been both harvested and subsequently planted after a crash in natural stands likely due to overharvesting (Buschmann et al. 2001). The artificial selection for tetrasporophytes has resulted in early stages of domestication (Valero et al. 2017) and loss of sexual reproduction (Guillemin et al. 2008). *Gracilaria vermiculophylla* (Ohmi) Papenfuss is a successful invader in many of the bays and estuaries of North America, northwestern Africa, and Europe (Krueger-Hadfield et al. 2017). The invasion success was likely facilitated by adaptive shifts in temperature and salinity tolerance (e.g., Sotka et al. 2018) and to biofoulers (e.g., Bonthond et al. 2020), as well as the ability to asexually fragment (Krueger-Hadfield et al. 2016). *Gracilaria caudata* J. Agardh can form dense stands in the intertidal zone (Plastino and Oliveira 1997) and has been subjected to intense harvesting pressure, leading to declines in native populations (Hayashi et al. 2014, see also Ayres-Ostrock et al. 2019). For this species, we re-analyzed the genome published by Flanagan et al. (2021). Finally, *Gracilaria gracilis* (Stackhouse) Steentoft, L.M. Irvine & Farnham is a long-lived species that inhabits tidepools along European coastlines. This species serves as model species to test hypotheses related to the evolution of sex (e.g., alternation of haploid and diploid phases in life cycles, Destombe et al. 1989, 1992, 1993, Hughes and Otto 1999; mating system and sexual selection, Richerd et al. 1993, Engel et al. 2002).

The availability of genomic and genetic resources for these four *Gracilaria* species should aid in our understanding of the evolutionary ecology of red algae in their dynamic environment, during invasions of new habitats, under cultivation practices, and in response to climate change. Moreover, these new resources will add to the existing genomic data and illuminate key processes in eukaryotic evolution. The Rhodoexplorer – Red Algal Genome Database currently includes the *Gracilaria* species discussed here but will include all the high-quality genomic resources available for the Rhodophyta (e.g., genomes, transcriptomes), thereby providing a unique resource for comparative analyses.

## RESULTS AND DISCUSSION

### Genome Assembly

Final genome assembly sizes, based on long and short read sequencing, ranged from 76 to 80 Mbp for *G. gracilis* and *G. chilensis,* respectively. In addition, we created a draft genome assembly based on the Illumina sequencing only for *G. caudata* (32 Mbp) and reassembled the genome of *G. vermiculophylla* (Flanagan et al. 2021) to a final 47 Mbp after bacterial contamination removal. The above genome assemblies were comparable to the genomes of *G. domingensis* (78 Mbp, Nakamura-Gouvea et al. 2022) and *G. changii* (36 Mbp, Ho et al. 2017). PacBio assemblies of *G. chilensis* and *G. gracilis* produced in this study (< 300 contigs per genome) are the most contiguous red macroalgal genomes presently available in public databases, apart from *G. vermiculophylla* and *P. yezoensis* where the addition of a HiC library enabled scaffolding nearly at the chromosome level (Wang et al. 2020, Flanagan et al. 2021). Despite the differences in assembly size, BUSCO scores were similar across the long read-sequenced *G. gracilis* and *G. chilensis*, and the more fragmented *G. caudata* genome, with 81.6 to 83.6% of conserved proteins present (Eukaryota_odb10, Manni et al. 2021, Simão et al. 2015; Table 1). The re-assembled genome of *G. vermiculophylla* contained 71.8% of the conserved proteins. Given the diversity of Rhodophyta and the lack of lineage-specific databases, these results are in the expected range. A recent study estimated the presence of conserved eukaryotic genes (Eukaryota_odb10) in red algal genomes at a median level of 69% (Hanschen et al. 2020).

Red algal genomes are repeat-rich, with half or more of their genomic sequence being constituted by repetitive elements, as reported previously for *Porphyra umbilicalis* (43.9%, Brawley et al. 2017), *Pyropia yezoensis* (48%, Wang et al. 2020) and *Chondrus crispus* (73%, Collen et al. 2013). In agreement with this general trend, between 45.7-66.2% of the *Gracilaria* genomes corresponded to repetitive elements (Figure 1 and Table 1).

**Fig. 1.**
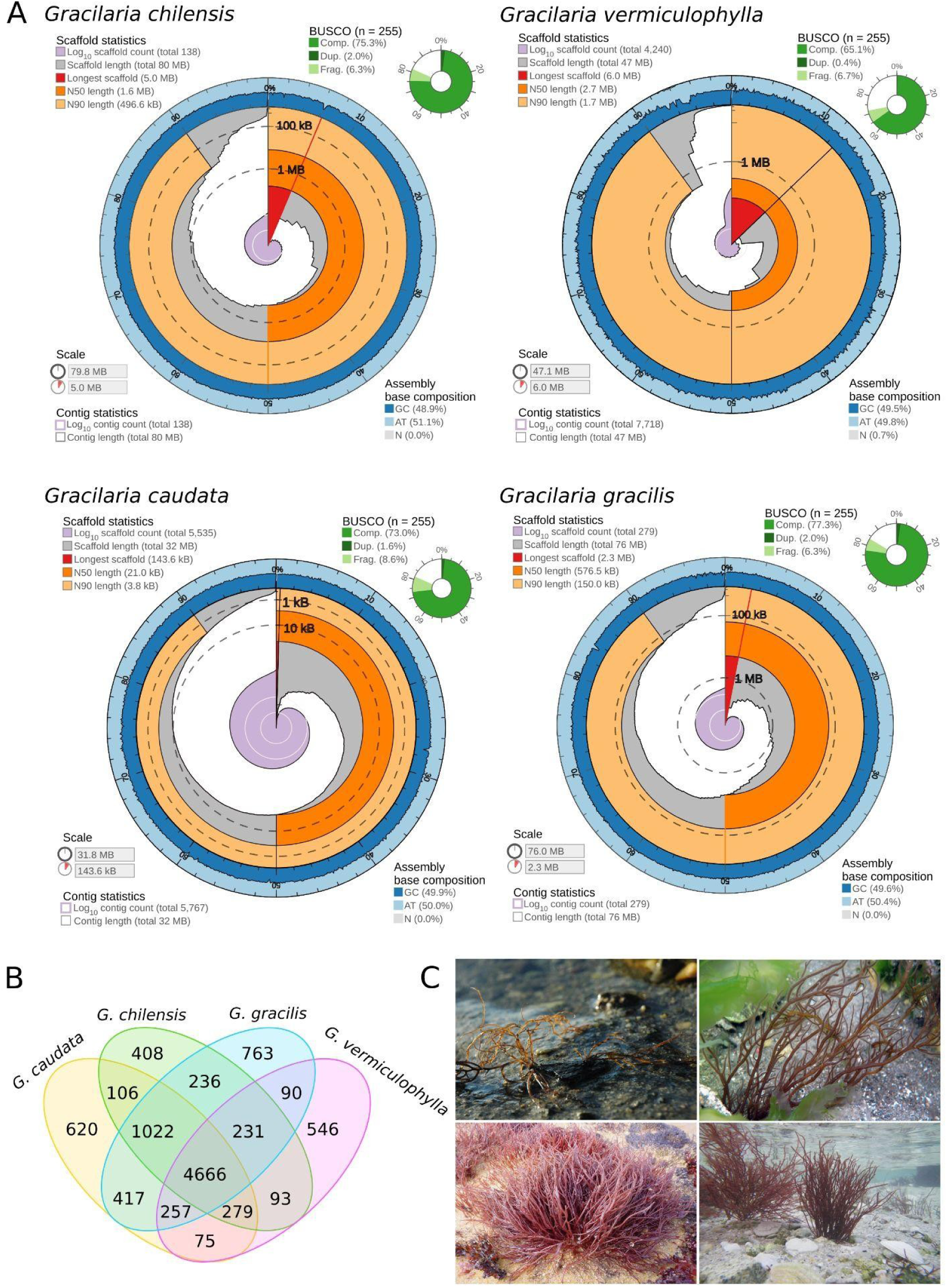
A) Genome assembly metrics of *Gracilaria chilensis* (top left), *Gracilaria vermiculophylla* (top right), *Gracilaria caudata* (bottom left) and *Gracilaria gracilis* (bottom right), (Challis 2017, https://github.com/rjchallis/assembly-stats). The inner radius (red) of the circular plot represents the length of the longest scaffold in the assembly and the proportion of the assembly that it represents. The cumulative number of scaffolds within a given percentage of the genome is plotted in light purple originating at the center of the plot. The N50 and N90 scaffold lengths are indicated by dark and light orange, respectively. Genome scaffolds are plotted in gray from the circumference and the length of segment at a given percentage indicates the cumulative percentage of the assembly that is contained within scaffolds of at least that length. The GC content is marked by the dark blue outer circle. Complete, fragmented and duplicated BUSCO genes are shown in green in the upper right corner. B) Venn diagram of shared and species-specific orthogroups and orphan genes among the four sequenced *G.* species. C) *G. chilensis* (top left), *G. vermiculophylla* (top right), *G. caudata* (bottom left) and *G. gracilis* (bottom right). Photo credit in order: M-L. Guillemin, S. Krueger-Hadfield, E. M. Plastino, C. Destombe.

**Table 1:**
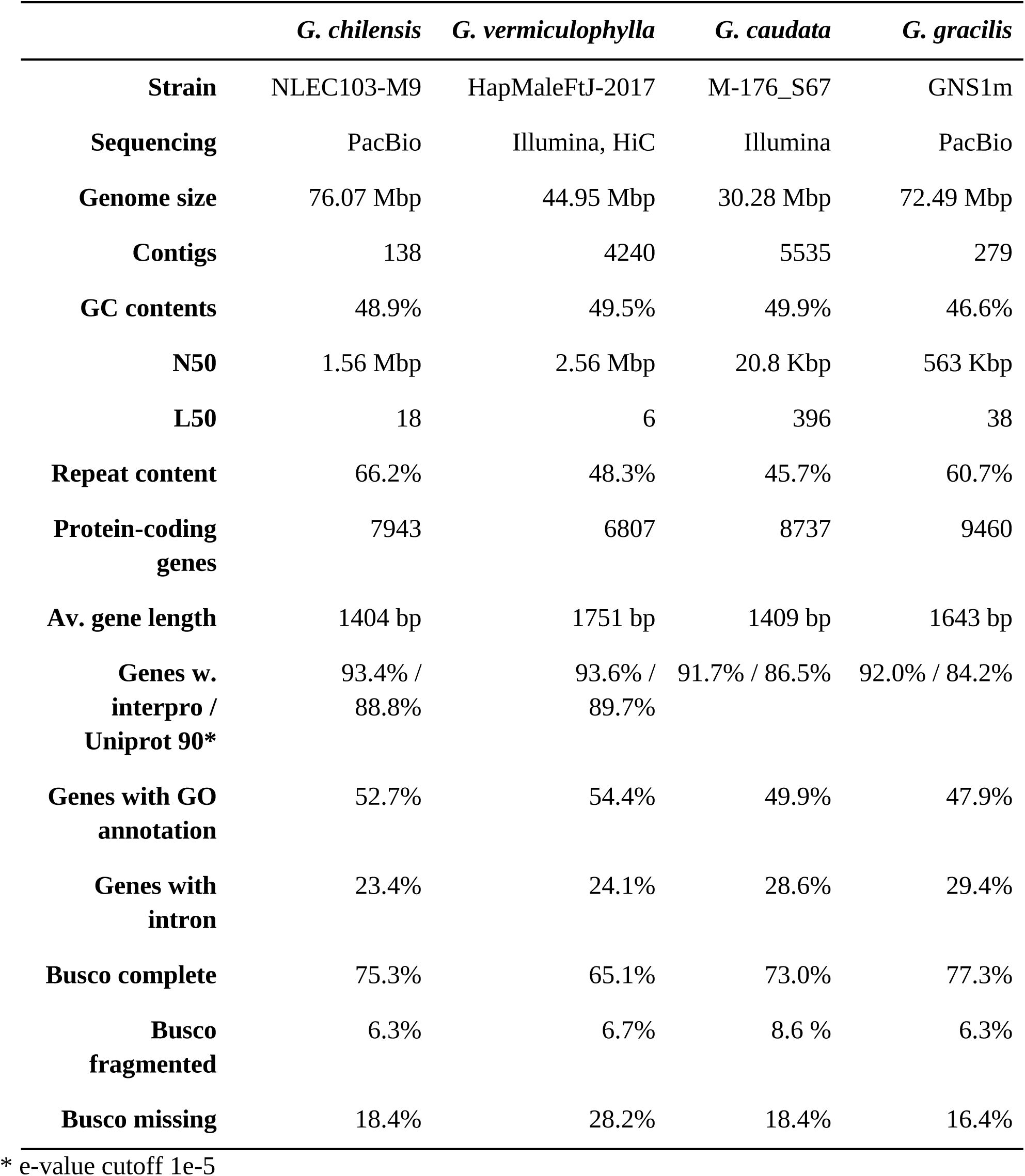
Assembly statistics.

### Gene prediction and Annotation

Gene prediction yielded a total of 8,042, 9,065 and 9,674 coding sequences for *G. chilensis*, *G. caudata* and *G.gracilis* (Table 1), which was comparable with other red macroalgal genomes, *Chondrus crispus* (9,815 genes, Collen et al. 2013) and *Gracilaria changii* genome (10,912 genes, Ho et al. 2022). In addition, we annotated the reassembled genome of *G. vermiculophylla*, which yielded fewer genes (7,048). Among these genes, 70.6-76.6% did not contain any introns, as typical for the compact genomes of red algae (Qiu et al. 2015). Most *Gracilaria* genes had homologous sequences in the Uniprot database (84.2-89.7%) and were annotated with at least one INTERPRO hit (91.7-93.6%). Between 47.9% and 54.4% of genes were associated with GO annotations.

Orthofinder analyses enabled us to identify 4,666 core groups of orthologous proteins present in all four of the sequenced genomes (Figure 1B) versus 408-620 orthogroups or orphan genes specific to only one of the sequenced species (Figure 1B). Among the species-specific sequences, the rate of GO annotation was lower than for the entire dataset, ranging from 12.7% for *G. chilensis* to 18.2% for *G. caudata*. The fact that the two species *G. caudata* and *G. gracilis* share more genes between them than with the phylogenetic group of *G. chilensis* and *G. vermiculophylla* was expected due to divergence between the two clades of *Gracilaria* species. Both the annotated and the unknown species-specific genes constitute attractive targets to study their role in adaptation and speciation.

### Rhodoexplorer – Red Algal Genome Database

In addition to depositing the raw reads and sequenced genome in a public repository (see Data Availability section), all four genomes were also integrated into the newly created Rhodoexplorer Red Algal Genome Database (https://rhodoexplorer.sb-roscoff.fr), hosted at the ABiMS bioinformatic platform. This platform will gradually include more red algal genomes in the future. The services provided include:

- Information about the sequenced strains, with links to external databases (NCBI, WoRMS, Algaebase)
- Assembly and annotation metrics
- Data downloads: genomic, genes and proteomic datasets, structural and functional annotations, orthology clusters, etc.
- A BLAST interface with a selection of red algal genomes, predicted and *de novo* assembled transcriptomes and proteomes.
- Visualization tools: a genome browser to visualize the predicted genes and the RNAseq data mapped on the genome and a web interface to visualize functional annotations and retrieve individual protein sequences.

## MATERIALS AND METHODS

### Sampling of the biological material

Adult female and male *Gracilaria* thalli, all bearing reproductive structures, used for the sequencing were collected from natural populations: *G. chilensis* in Lenca (Chile, −41.607, −-72.692), *G. vermiculophylla* in Charleston, SC (USA, 32.752, −-79.900), *G. caudata* in Paracuru, CE (Brazil, −-3.399, −-39.012), and *G. gracilis* in Cape Gris-Nez (France, 50.872, 1.584). *Gracilaria caudata* and *G. chilensis* were maintained as clonal, unialgal cultures under laboratory conditions prior to nucleic acid extractions (see *Culture conditions*). Field-collected *G. gracilis* and *G. vermiculophylla* thalli were transported to the laboratory, examined under a microscope, and cleaned of contaminants. If visible, cystocarps were excised prior to preservation of the thalli at −-80℃ before further use. Table S2 provides details of the *Gracilaria* species used in this study.

### Culture conditions

Cultures were initiated either from lab crosses or from tetraspores released by field-collected tetrasporophytes. *Gracilaria caudata* was grown in the modified von Stosch nutrient solution (Ursi and Plastino 2001) diluted to 25% in seawater (32 psu), with weekly renewals. The algae were kept in culture chambers at 25°C under fluorescent illumination of 70 μmol.m-2.s-1 14h photoperiod, following previously established optimal growth conditions (Yokoya and Oliveira 1992a,b). *Gracilaria chilensis* was grown in Provasoli medium (McLachlan 1973), changed weekly during the first two months and twice a week thereafter. Cultures were kept at 13°C under 40-60 μmol.m-2.s-1 of light with 12h day length.

### Nucleic acid extraction, library preparation, and sequencing

Genomic DNA was extracted from mature male gametophytes using DNeasy PowerPlant Pro Kit for *G. caudata* or an in-house protocol based on Faugeron et al. (2001) for *G. chilensis* and *G. gracilis*. The concentration and purity of DNA were measured with NanoDrop and Qubit before sequencing on an Illumina HiSeq 2500 (125 bp PE reads for *G. chilensis* and *G. gracilis*; 100bp PE reads for *G. caudata*) or PacBio Sequel II with sheared gDNA large insert library (*G. gracilis* and *G. chilensis*) (Table S2).

For genome annotation, total RNA was extracted from mature thalli of male and female gametophytes of *G. chilensis* (2 males and 2 females), *G. caudata* (4 males and 4 females), and *G. gracilis* (1 male and 1 female) using the Rneasy Mini Plant Kit (Qiagen) following the manufacturer’s instructions. Total RNA was extracted from *G. vermiculophylla* (4 males and 4 females) using the Macherey Nagel Nucleospin RNA for Plant kit following the manufacturer’s instructions. Paired-end 150bp Illumina reads were generated with Illumina HiSeq 2500 Table S2).

### Genome assembly

De novo genome assemblies for *G. gracilis* and *G. chilensis* were generated based on 203-fold and 116-fold coverage of PacBio long reads, respectively. Briefly, bacterial sequences were removed from raw data (subreads) using Blobtools v1.1.1 (Laetsch and Blaxter 2017). For each species, two independant assemblies were generated using CANU (Koren et al., 2017) and FLYE (Kolmogorov et al., 2019). Based on congruity (QUAST v.5.0.2 – Mikheenko, et al., 2018) and BUSCO score (Simão FA, et al. 2015) the best assemby was kept and polished using three iterations of RACON v.1.4.20. Finally, PacBio sequencing error were corrected using 150bp paired-end Illumina reads with PILON v.1.23 software (Walker et al. 2014). The draft genome assembly of *G. caudata*. was generated using 171-fold coverage of 150bp paired-end Illumina reads only. First, a meta-genome was produced using metaSPAdes v3.12.0 (Nurk et al. 2017) and bacterails contigs were detected using Blobtools. Reads corresponding to eukaryotic contigs were then assembled using SPAdes v3.12.0 (Bankevich et al. 2012). Quality of all *de novo* genome assemblies was assessed with QUAST and DNAseq remapping for congruity and BUSCO and RNAseq mapping for completness.

For *G. vermiculophylla*, we updated the existing chromosome-scale genome assembly (Flanagan et al. 2021) by reassembling the Illumina reads using SPAdes v 3.12.0 (Bankevich et al. 2012) and scaffolding with Hi-C libraries, following the Dovetail Genomics proprietary pipeline (Elbers et al. 2019). This process ameliorated the genome continuity (N50 increased from 2.06Mb to 2.68Mb) and completeness (BUSCO score increased from 57,6% to 65,9% of complete genes using the Eukaryota_odb10 dataset).

We used Blobtools v1.1.1 (Laetsch and Blaxter 2017) with maximal accuracy settings to validate the quality of the four *Gracilaria* genome assemblies and identify potential bacterial contaminations. In brief, DNAseq reads of each species were first mapped to their corresponding reference genome using HISAT2 v 2.2.1 (Kim et al. 2019). Next, BAM coverage files produced by HISAT2, Diamond blastx v2.0.11 (Buchfink et al. 2015, 2021) hit-file against non-redundant protein sequences archive from NCBI (-sensitive, –max-target-seqs 1, -e-value 1e − 20), and Blast v2.12.0 (Camacho et al. 2009) output against nucleotide archive from NCBI (-max_target_seqs 10 -max_hsps 1 -evalue 1e − 20) were used as input for Blobtools. Genomic scaffolds classified as bacterial or with a coverage of less than 1 (sum of coverages for each sequence across all coverage files) were removed from the assembly. Genome assembly completeness was assessed using BUSCO scores with the eukaryotic data set (Eukaryota_odb10, Simão et al. 2015, Manni et al. 2021).

Chloroplastic and mitochondrial genomes of each species were reconstructed from Illumina raw reads using NovoPlasty (Dierckxsens et al. 2016) throught the European Galaxy web portal (https://usegalaxy.eu/). Annotation of those *de novo* organellar genomes were done using the GeSeq web tool (Tillich et al. 2017 – https://chlorobox.mpimp-golm.mpg.de/geseq.html). Public sequences from *Gracilaria caudata* voucher SPF:57390 (NC_039146, NC_039139), *Gracilaria chilensis* voucher CNU050183 (KP728466, KT266788), *Gracilaria gracilis* voucher SPF:55734 (NC_039141, NC_039148) and *Gracilaria vermiculophylla* (MN853882, MH396022) were retrieved from NCBI and used as seeds and references for both assembly and annotation.

### Genome annotation

Each reference genome was first masked using RepeatMasker v4.0.9 (Smit et al. 2015) with Dfam v3.0 database (Wheeler et al. 2013) and a customized repeat library produced from concatenated outputs of RepeatScout v1.0.6 (Price et al. 2005) and TransposonPSI v1.0.0 (Hass 2007-2011). Initial quality assessment of the RNA-Seq reads was performed with FastQC v0.11.9 (Andrews et al. 2010), and reads were trimmed using Trimmomatic v0.39 (TRAILING:3 SLIDINGWINDOW:4:15 MINLEN:50; Bolger et al. 2014). Clean reads were mapped to the reference genome assembly using HISAT2 v 2.2.1 (Kim et al. 2019). The resulting alignment files were used to annotate protein-coding genes with BRAKER2 v2.1.6 pipeline (Bruna et al. 2021). Functional annotation of the reference transcriptomes was performed using eggNOG-mapper (Huerta-Cepas et al. 2019, Cantalapiedra et al. 2021).

All code used for genomes assembly and annotation is available on the Gitpage dedicated to the genome database project https://abims-sbr.gitlab.io/rhodoexplorer/doc/data_process/.

### Rhodoexplorer – Red Algal Genome Database

The main web portal (https://rhodoexplorer.sb-roscoff.fr) has been implemented using the Python web framework Django, with data stored in a relational database (PostgreSQL).

For each red algal species, an integrated environment of visualization tools has been deployed based on the Galaxy Genome Annotation (GGA) project (Bretaudeau et al. 2019). Each GGA environment deployed for the Rhodoexplorer genome database includes: Chado – a PostgreSQL relational database schema for storing biological data (Mungall et al. 2007); JBrowse – a web-based genome browser (Buels et al. 2016); Tripal – a Drupal-based application for creating biological websites (Sanderson et al. 2013); Elasticsearch – a distributed, free and open search and analytics engine for all types of data (https://www.elastic.co/products/elasticsearch); Galaxy – a browser accessible workbench for scientific computing used as a data loading orchestrator for administrators (The Galaxy Community 2022). To facilitate the deployment and the administration of the GGA service, a set of Python tools has been developed (http://gitlab.sb-roscoff.fr/abims/e-infra/gga_load_data) allowing mass deployment of Docker containers and automated data loading through Galaxy with the Bioblend API (Sloggett et al 2013).

The BLAST interface (https://blast.sb-roscoff.fr/rhodoexplorer/) includes an implementation of the BLAST algorithm using SequenceServer (Priyam et al. 2019) graphical.

The documentation website for navigating the platform web portal and resources (https://abims-sbr.gitlab.io/rhodoexplorer/doc/) is published from a GitLab repository, with Pages and MkDocs, a static site generator.

The entire informatic infrastructure is deployed and maintained on the ABiMS Bioinformatics platform of the Roscoff Biological Station, part of the national infrastructure French Bioinformatic Institute.

## SUPPLEMENTARY MATERIAL

Supplementary Figure S1: Life cycle of *Gracilaria*.

Supplementary Table S1: Available red algal genomic resources.

Supplementary Table S2: Species used in this study.

## ACKNOWLEDGEMENTS

This project was supported by start-up funds from the College of Arts and Sciences at the University of Alabama at Birmingham to SAKH; ANID NCN2021-033 and FONDECYT 1221456 and 1221477 to MLG, JB and SF; the International Research Networks DEBMA “Diversity, Evolution and Biotechnology of Marine Algae” (CNRS GDRI 0803) and DABMA “Diversity, Adaptation, and Biotechnology of Marine Algae” (CNRS IRN 00022); the ERC (grant number 864038 to SMC); and the ANR project IDEALG (ANR-10-BTBR-04, “Investissements d’Avenir, Biotechnologies-Bioressources”). We are grateful to the Roscoff Bioinformatics platform ABiMS (http://abims.sb-roscoff.fr), part of the Institut Français de Bioinformatique (ANR-11-INBS-0013) and BioGenouest network, and the Max Planck Institute for Biology Tubingen for providing computational resources. We also wish to thank Kristy Hill-Spanik, Rosário Petti, and Vivian Viana for field and technical support.

## DATA AVAILABILITY

Sequencing data has been deposited in the SRA database under BioProjects PRJNA936482, PRJNA931233, PRJNA938301, PRJNA938403. The accession numbers for the raw sequence data are provided in Table S2.

*Gracilaria chilensis, Gracilaria gracilis* and *Gracilaria caudata* Whole Genome Shotgun project have been deposited at DDBJ/ENA/GenBank under the accessions JARGXX000000000, JARGSG000000000 and (awaiting accession number), respectively. The versions described in this paper are versions JARGXX010000000, JARGSG010000000 and (awaiting accession number). *Gracilaria vermiculophylla* updated assembly has been deposited under (awaiting accession number).

## Footnotes

1 There is controversy over the systematics of Gracilaria Greville, but for the purposes of this paper, we consider the four species as belonging to the genus Gracilaria (sensu Lyra, et al. 2021, Guiry and Guiry 2022).

## SUPPLEMENTARY TABLES AND FIGURES

**Fig. S1.**
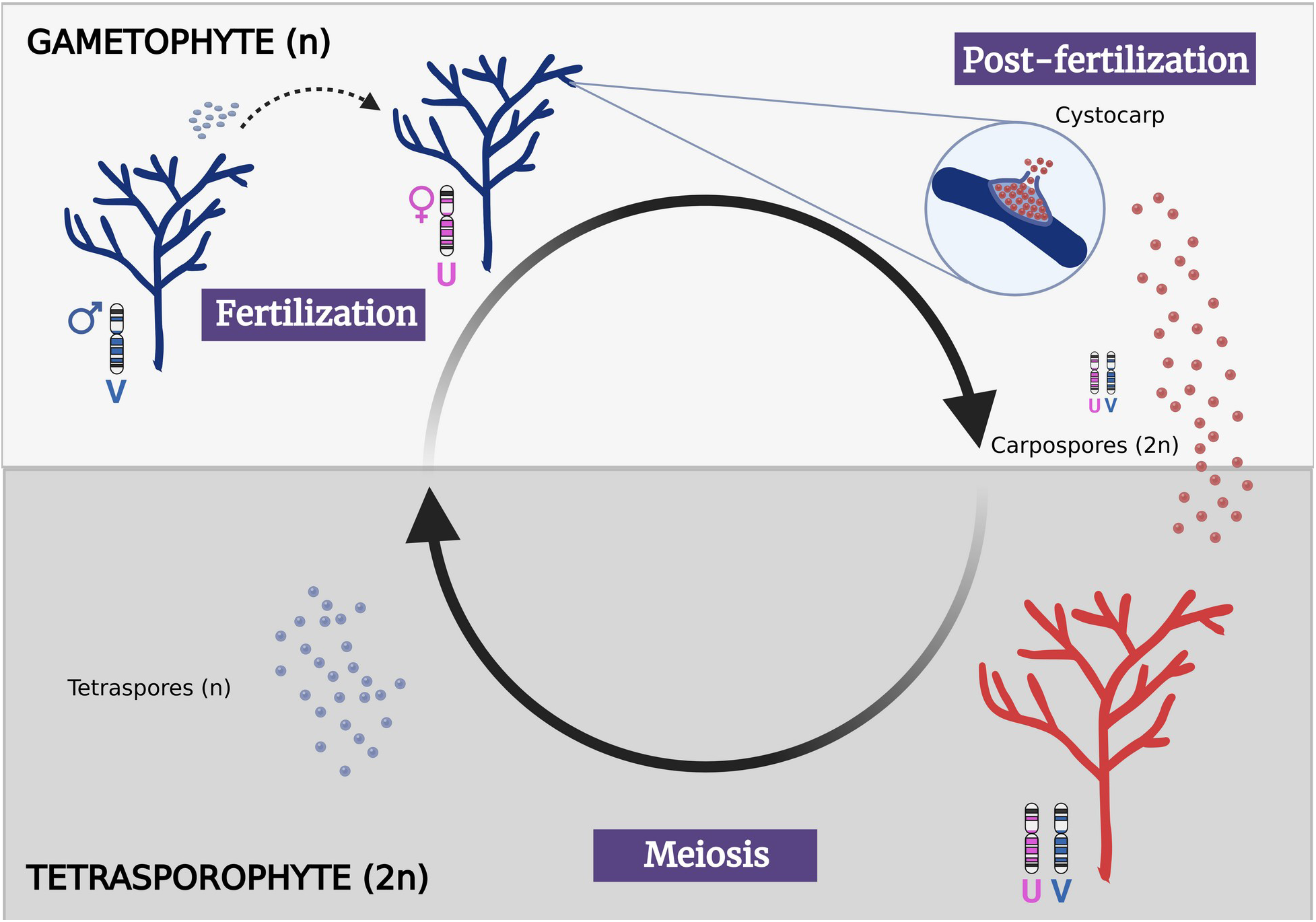
Life cycle of *Gracilaria.* The life cycle consists of an alternation between haploid dioecious gametophytes and a diploid tetrasporophyte. The tetrasporophyte produces meiospores through meiosis, which develop as gametophytes after release. The sex of the gametophytes is determined by haploid sex chromosomes (UV system). Spores that receive the V sex chromosome develop as male gametophytes whereas spores that carry U chromosome will produce female gametophytes. After fertilization, the zygote develops within the carposporophyte on the female gametophyte and is mitotically amplified—producing thousands of diploid carpospores that after release will give rise to tetrasporophytes.

**Supplementary Table S1:** Available red algal wholes genome sequences. M=multicellular, U=unicellular.

| Species | Order | N50 | U/<br>M | Citation |
| --- | --- | --- | --- | --- |
| *Chondrus crispus* | Gigartinales | 250k<br>b | M | https://doi.org/10.1073/pnas.1221259110 |
| *Galdieria sulphuraria* | Cyanidiales | 230k<br>b | U | https://doi.org/10.7554/eLife.45017 |
| *Galdieria phlegrea* | Cyanidiales | 201k<br>b | U | https://doi.org/10.7554/eLife.45017 |
| *Gracilaria changii* | Gracilariales | 17kb | M | https://doi.org/10.1016/j.ygeno.2017.09.003 |
| *Gracilaria domingensis* | Gracilariales | 189k<br>b | M | https://doi.org/10.1111/jpy.13238 |
| *Gracilaria vermiculophylla* | Gracilariales | 2Mb | M | https://doi.org/10.1111/mec.15854 |
| *Gracilariopsis chorda* | Gracilariales | 220k<br>b | M | https://doi.org/10.1093/molbev/msy081 |
| *Gracilariopsis lemaneiformis* | Gracilariales | 35kb | M | https://doi.org/10.1186/s12870-018-1309-2 |
| *Calliarthron tuberculosum* | Corallinales | n/a | M | https://doi.org/10.1016/j.cub.2011.01.037 |
| *Porphyridium purpureum* | Porphyridiales | 20kb | U | https://doi.org/10.1038/ncomms2931 |
| *Porphyra umbilicalis* | Bangiales | 202k<br>b | M | https://doi.org/10.1073/pnas.1703088114 |
| *Neoporphyra haitanensis* | Bangiales | 650k<br>b | M | https://doi.org/10.1093/molbev/msab315 |
| *Neopyropia yezoensis* | Bangiales | 34M<br>b | M | https://doi.org/10.1038/s41467-020-17689-1 |
| *Kappaphycus alvarezii* | Gigartinales | 849k<br>b | M | https://doi.org/10.1101/2020.02.15.950402 |
| *Asparagopsis taxiformis* | Bonnemaisoniales | 2kb | M | https://doi.org/10.1021/acschembio.0c00299 |
| *Cyanidium caldarium* | Cyanidiales | 13kb | U | https://www.ncbi.nlm.nih.gov/genome/7354* |
| *Cyanidiococcus yangmingshanensis* | Cyanidiales | 653kb | U | https://doi.org/10.1111/jpy.13056 |
| *Cyanidioschyzon merolae* | Cyanidioschyzonales | 846kb | U | https://doi.org/10.1186/1741-7007-5-28<br>https://doi.org/10.1038/nature02398 |

\* no publication associated n/a data no longer accessible

**Supplementary Table S2:**
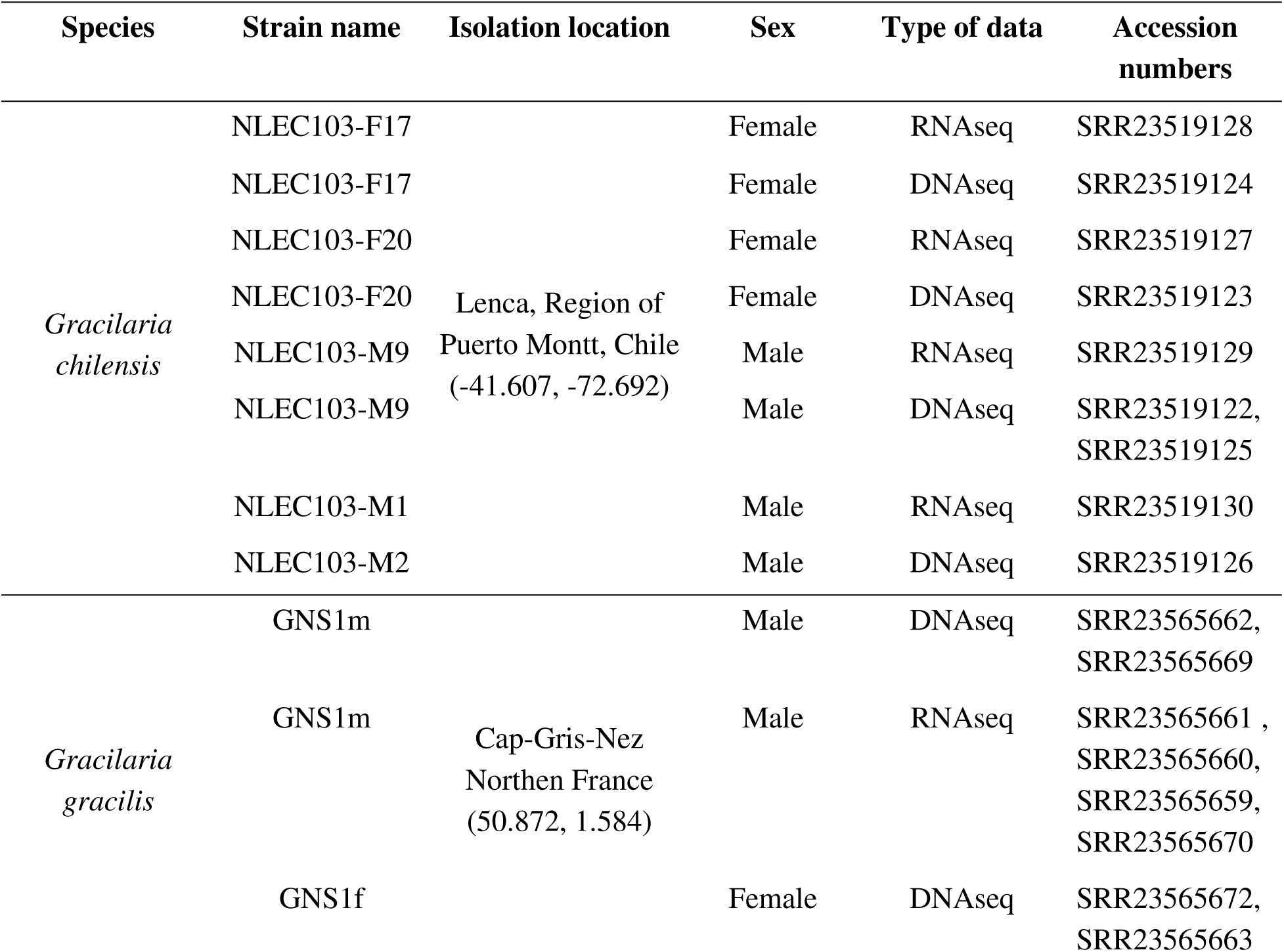
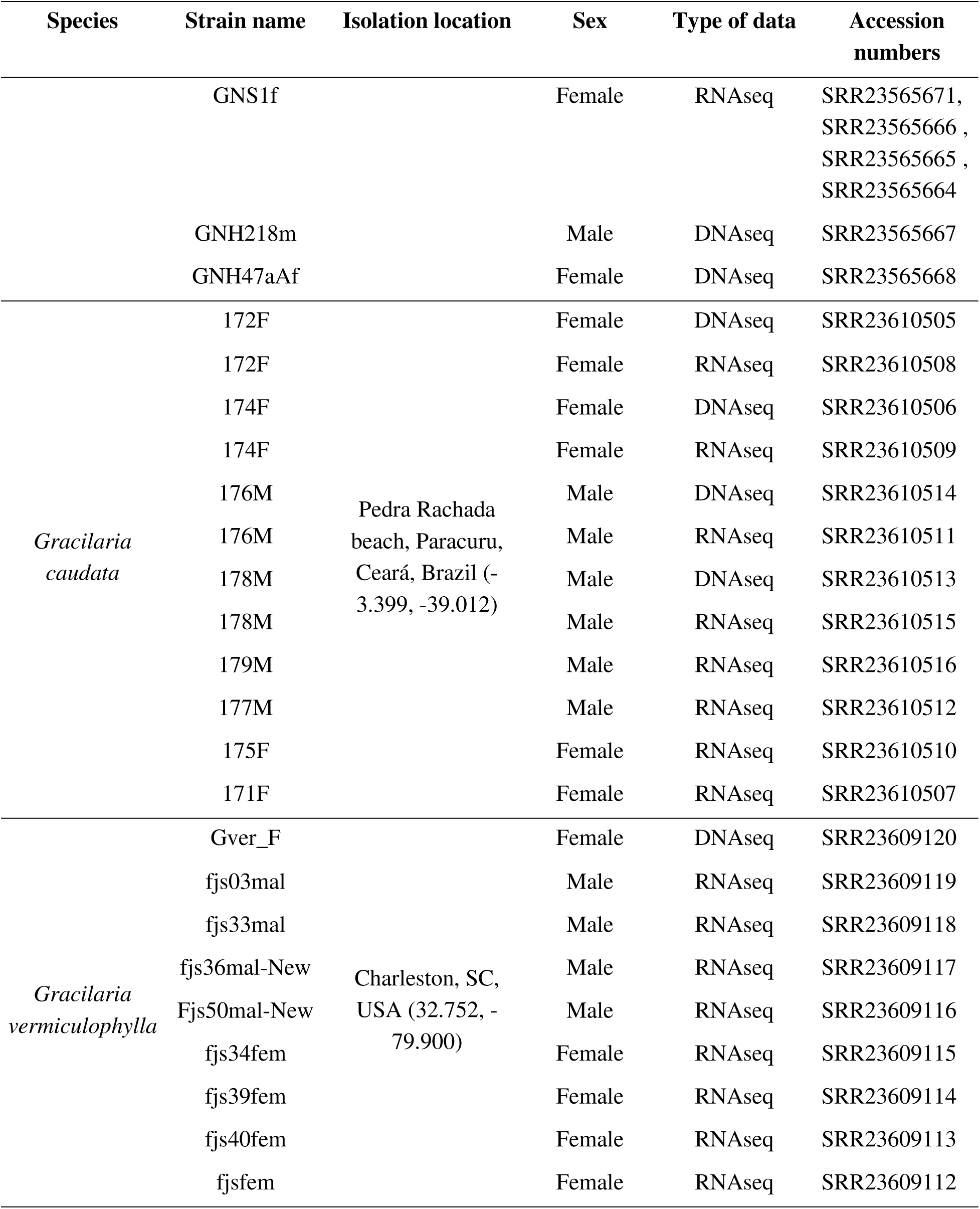
Species used in this study

